# Boost the resilience of Protected Areas to shocks by reducing their dependency on tourism

**DOI:** 10.1101/2022.11.21.517412

**Authors:** F. Ollier D. Andrianambinina, Derek Schuurman, Mamy A. Rakotoarijaona, Chantal N. Razanajovy, Honorath M. Ramparany, Serge C. Rafanoharana, H. Andry Rasamuel, Kevin D. Faragher, Patrick O. Waeber, Lucienne Wilmé

## Abstract

Nature-based or ecotourism is widely considered a strong mechanism for the sustainable funding of protected areas (PAs). Implemented during the 1990s in Madagascar, nature-based tourism experienced positive growth over the last 30 years with increasing numbers of visits to the parks and reserve safeguarding the endemic biodiversity. Revenue earned from entrance fees to the network of PAs managed by Madagascar National Parks has never been sufficient to finance their management. Political crises and the COVID-19 pandemic in particular, have highlighted the risks for park managers of relying on such earnings when they covered just 1 % of the funding required in 2021. Alternative mechanisms of funding are analysed for all Madagascar’s PAs in order to facilitate sustainable conservation of the localities and protection of the island’s biodiversity.

## Introduction

For a long time, nature-based tourism—for our purposes being defined as tourism centered around the appreciation of wild animals in their natural habitats—has been regarded as a mechanism which contributes to the successful conservation of protected areas (PAs) by increasing their visibility and in so doing, attracting political attention, encouraging financial support, raising awareness of nature and ultimately, safeguarding biodiversity. It is thus also held to play a pivotal role in financing conservation actions [1].

Studies have revealed that nature-based tourism tends to bring more visitors to those PAs with the highest levels of biodiversity [2–4]; to those which have been established longer [5]; those that are larger in size [6,7], and those which are more readily accessible from urban areas [8]. Other factors which have been shown to influence visitor numbers include climate and weather [9,10] as well as elevation [8] It has been found that fewer people visit the more remote PAs, while PAs in high income countries tend to receive more visitors [11,12].

Mainland Africa is renowned for its trans-boundary peace parks and a host of other protected areas. Millions of tourists visit the classic African safari destinations such as Kenya, Tanzania, Botswana, Zambia, and South Africa, to observe larger mammals, especially the so-called ‘Big Five’ (African elephant, Cape buffalo, African lion, Leopard and Rhinoceros). While the dark spectre of wildlife crime perpetually looms in the background, threatening to undermine nature-based tourism, sub-Saharan Africa’s safari industry is estimated to have generated US$ 12.4 billion of annual revenues in 2019 [13] while the continent’s wildlife and nature-based tourism industry as a whole, is estimated to generate in excess of US$ 29 billion annually [14].

Apart from safari tourism, wildlife watching tourism in Africa includes products such as Gorilla and Chimpanzee tracking in Central Africa and also on the theme of primate watching experiences, just across the Mozambique Channel, Madagascar offers Lemur watching as its primary unique selling point. While Madagascar lacks the ‘Big Five’ or great apes, the island does have an exceptionally high degree of endemism in its biodiversity, including the >100 species of lemurs which only occur there. These charismatic non-human primates range in size from the monogamous, blue-eyed and ape-like Indri (*Indri indri*) which weighs in at some 7 kg, to the diminutive Madame Berthe’s mouse-lemur (*Microcebus berthae*) which tips the scale at only 0.050 kg. The contrasting and fluffy coats of many of the larger lemur species, add to their appeal and popularity [15]. Of the world’s eight baobab species, six occur only in Madagascar [16].

While Madagascar is less than a sixth the size of the Congo basin in Central Africa, it has more plant species than the latter (14 thousand VS. 10 thousand) and the rate of endemism is 80 % for Madagascar plants as compared to 30 % for the Congo basin. In Madagascar, rates of endemism among its faunal groups vary from 50 % for the birds, to almost 100 % for reptiles and amphibians [17]. With about 5000 km of shoreline, the island is mostly pantropical and contains a diversity of forests from humid and sub-humid in the East and Northeast, to dry deciduous and dry ‘spiny bush’ or sub-arid thorn thickets in the West and Southwest [17–19]. Throughout its coastal waters, Madagascar provides breeding and calving habitat for humpback whales (*Megaptera novaeangliae*) from June to September before migrating back to polar waters where they feed [20,21].

The island’s terrestrial biodiversity is mostly concentrated in forests and to a lesser extent in wetlands, with a sprinkling of endemic taxa inhabiting the open landscapes dominated by grasslands [17,19,22,23]. The largest and most undisturbed forests have been protected as Strict Nature Reserves since the first half of the 20th century. The protected areas network was created to safeguard maximal biodiversity and in so doing, cover most of the island’s ecosystems, including the marine environment. A principle aspect has always been to ensure connectivity through corridors. Of the 123 PAs, 96 are terrestrial; 9 marine and 18 include both terrestrial and marine ecosystems. Nine of the eleven Strict Nature Reserves (IUCN Ia) have been turned into National Parks (IUCN II) to accommodate nature-based tourism. Forty-three PAs, with IUCN categories I, II, and IV [24], are managed by MNP—Madagascar National Parks, a parastatal organization.

The 1990s saw the emergence of the nature-based tourism concept after the World Bank and International Development Bank—motivated by global rainforest losses—had discontinued loans to mass tourism organizations, reinstating them in 1990 under the nature-based tourism banner [25]. It was also during that period when nature-based tourism began to develop in Madagascar [26]. Annual numbers of inbound visitors to Madagascar increased slowly from approximately 100,000 in the mid-1990s to almost 500,000 in 2019, and this resulted in a total revenue just short of US$ 1 billion [27]. Clearly therefore, tourism assumed a significant position within the total market value of final goods and services produced in Madagascar, i.e., around 6 % Gross National Product (GNP) in 2019. While the statistics appear encouraging, it should be pointed out that before Madagascar’s PA network was expanded in mid-2010, the collective earnings from tourism entrance fees were significantly lower than the maintenance costs for effective PA management were [28]—and that this is still the case.

The onset of the COVID-19 pandemic brought the world to a quasi-halt in March 2020. The international aviation industry, which had registered almost 40 million flights in 2019 globally, recorded just below 17 million and 19 million flights for 2020 and 2021, respectively [29]. Tourism is one of the sectors that suffered most extensively as a result of the pandemic. In March 2020, Madagascar closed its borders, only reopening them again in April 2022. The Protected Areas were off limits to visitors from March to July 2020.

Despite trends indicating increases through the last 20 years, visitor numbers to Madagascar’s PAs are still too low for the collective entry fee receipts to contribute sufficiently to PA management costs. We therefore hypothesize that tourism on its own is still not paying for conservation. And we argue that new financial solutions for PA management need to be formulated, especially since—as was demonstrated by the COVID-19 pandemic – tourism is prone to external shocks which negatively impact park entry-based revenues.

We examine protected areas that potentially harbour higher biodiversity, and which therefore are assumed to be more alluring to visitors [2]. Our focus is on PAs that are managed for tourism, allowing for comparisons over differing time periods.

## Methodology

For this study we selected terrestrial protected areas (PAs) with IUCN Ia (Strict Nature Reserve); IUCN II (National Parks) managed for ecosystem protection and recreation) and IUCN IV status (Habitat / Species Management Areas, managed for conservation through management interventions) [30] (Fig 1). We have not included the other IUCN categories—firstly, IUCN III (Natural Monuments, managed for special natural features), of which only two examples, both small, exist—nor have we examined IUCN V and VI as those are created for multiple usage and as recently as the mid-2010s [28,31]. As such, they have a limited history.

**Fig 1.**
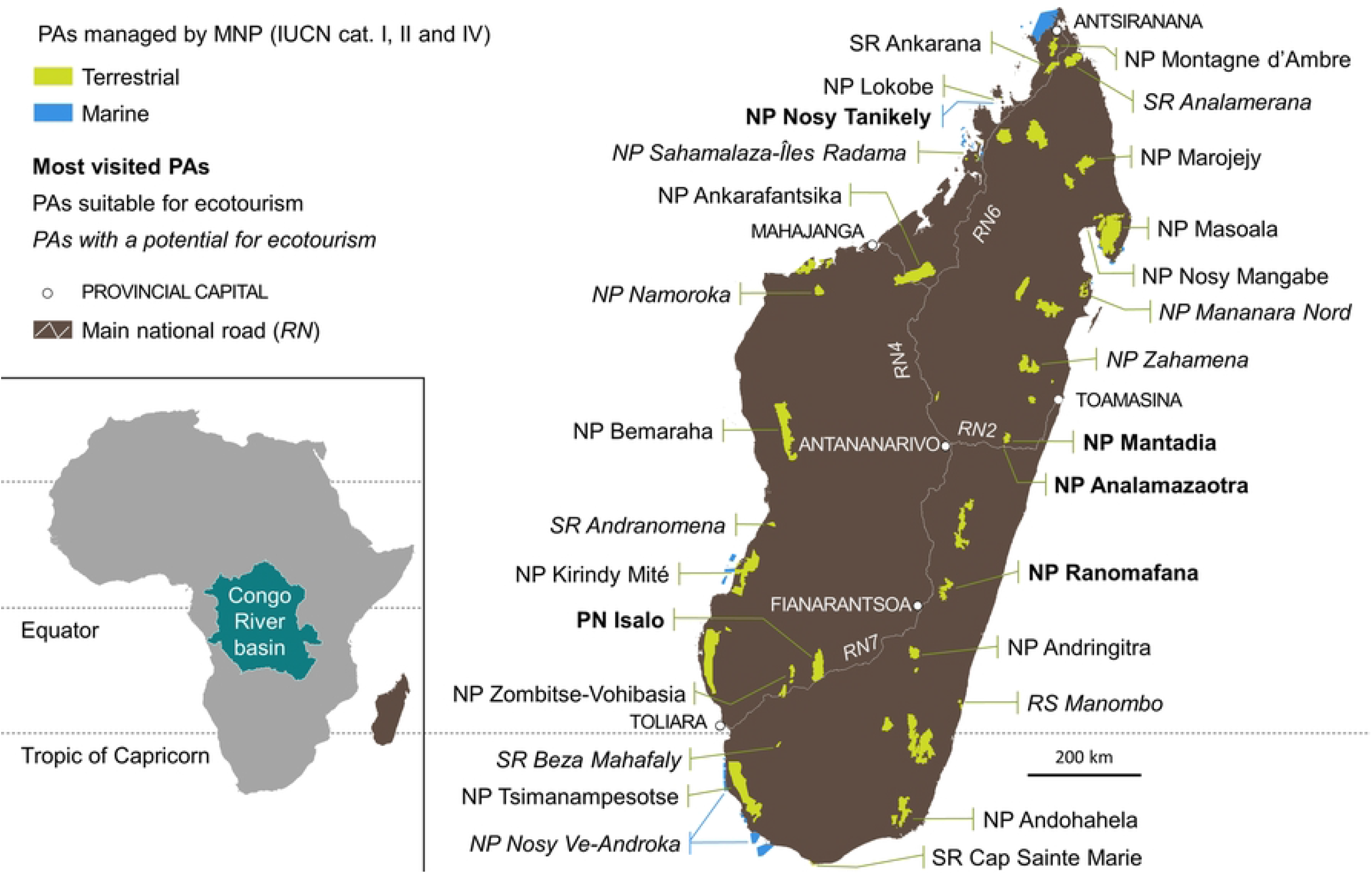
Protected Areas Madagascar. Protected areas in IUCN categories I, II and IV managed by Madagascar National Parks (MNP) and their importance for the implementation of nature-based tourism.

Another filter we applied in the selection of PAs is the management structure. We consider only PAs managed by MNP, because they have a centralized and standardized governance approach which also facilitates access to data. The PAs considered, account for 43 parks and reserves managed by MNP: in 20, nature-based tourism has been implemented, in 10 there is potential for the development of this activity while the remaining 13, are not considered suitable for the development of nature-based tourism (Table A in S1 File).

We collected and collated annual and monthly visitor data over the 30-years period from 1992 to 2021. These data reveal the MNP income for PA management from tourism. To assess the net contribution of tourism to PA management, we also compiled and collated MNP’s costs of managing the network of the PAs they oversee, considering elements such as staff, external services, consumables, and infrastructure / equipment, as well as their sources of funding from 2017–2021. We focused on these years because of changes made to the entrance fee structure at the end of 2016. We gathered the total number of tickets sold for each Protected Area from January 1992 to December 2021 (Table B in S1 File). Note that the number of tickets sold does not reflect the number of individual visitors: there is no specific ID associated with any particular ticket, so an individual visitor can have several PA entries for different dates. All data are from MNP.

To ascertain the impact of the COVID-19 pandemic on the revenue generated by park entrance fees, we calculated a trend line for PA visits by focusing on the number of tickets sold during years that were not affected by major crises, i.e., the periods from 1992 to 2001; 2003 to 2009, 2011 to 2019 for terrestrial PAs, as well as the years 2011 to 2019 for marine PAs. The ten years from 1992 to 2001 were politically stable, while the year 2002 was marked by a political crisis in January following the presidential election during which Marc Ravalomanana opposed Didier Ratsiraka, and the years 2009 and 2010 were strongly impacted by the January 2009 coup d’état after presidential elections, when Marc Ravalomanana and Andry Rajoelina were in opposition to one-another, and which resulted in a prolonged period of political instability. In the process of calculating the increase in the numbers of PA visits since 1992, we consider 2019 as being a ‘normal’ year, even though 2019 followed two political crises, a period of insecurity and sporadic failures in the provision of services to visitors. We estimated losses due to shortfalls in sales of entrance permits for 2020 and 2021 based on the trends calculated over the mentioned 30 years and the average distribution of visits per PA for the period 2010–2019, including the entire network of PAs where there is nature-based tourism. We took into account age and nationality of visitors to the different PAs, after the establishment of the new entrance fees, in order to analyse the different rates of entrance fees the period 2017 and 2019 in marine PAs and terrestrial PAs, i.e., after the establishment of the new entrance fees, to properly consider the different rates of entrance fees (Table C in S1 File).

For data interpretation and validation, we conducted a survey of park managers from the main PAs managed by MNP who participated in a workshop in Antananarivo during February 2022. The 17 park managers were questioned about how they would explain a shortage or a change in tourist numbers to their parks and they were asked to elaborate on the socio-economic and/or political context pertaining to specific years in their regions.

Investigating alternative funding sources for park management, we also consulted with FAPBM (*Fondation pour les Aires Protégées et la Biodiversité de Madagascar*, Foundation for the protected areas and the biodiversity of Madagascar) to detail the mechanisms and contribution of funds towards protected area management and conservation in Madagascar.

## Results

The number of tickets sold to enter parks and reserves from 1992 to 2019 increases over the years, with significant declines for certain years being followed by gradual recoveries (Fig 2). Considering the number of tickets sold for years not affected by major crises, i.e., the periods from 1992 to 2001; 2003 to 2009 and 2011 to 2019 for terrestrial PAs, as well as the years 2011 to 2019 for marine PAs, the increase in the number of tickets sold to enter parks and reserves is linear (Fig 2).

**Fig 2.**
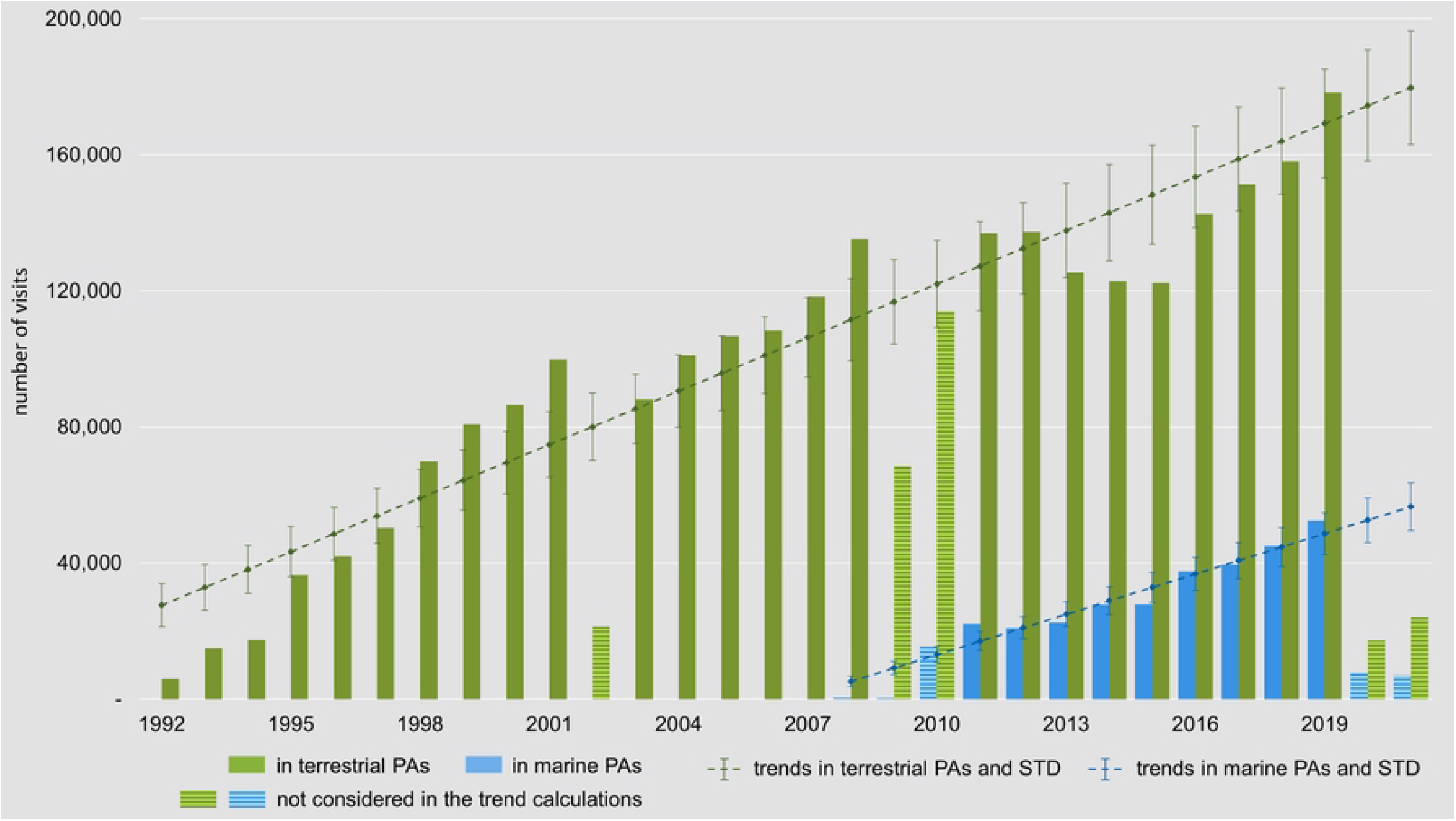
Park entries. Total number of visits in parks and reserves managed by MNP and trend lines calculated for both terrestrial protected areas and marine protected areas. (Trend line for terrestrial PAs: number of visits = 5,246 * (year - 1991) + 22,354 (r^2^=0.90); for marine PAs: number of visits = 3,952 * (year - 2010) + 13,090 (r2=0.93); 2002, 2009–2010, and 2020–2021 are not considered in the estimation of the trends as they are years of political instability; despite the financial crisis end of 2008, the year 2008 has been considered as “normal” given that visitors to Madagascar plan their visit ahead of time, and number of visitors to PAs has not declined end of 2008)

From 2016 to 2019, the entrance fees and secondary or additional incomes from tourism covered only approximately 35–40 % of the conservation management cost, compared to less than 20 % during years 2013–2015. Over a 26-year period from 1995 to 2020, a total of 5,717,100 tourists was reported for Madagascar, generating an estimated revenue of US$ 10,5 billion for the period 1995–2020 [29,32]. During this time, 2,938,736 tickets to enter PAs were sold (Fig 3).

**Fig 3.**
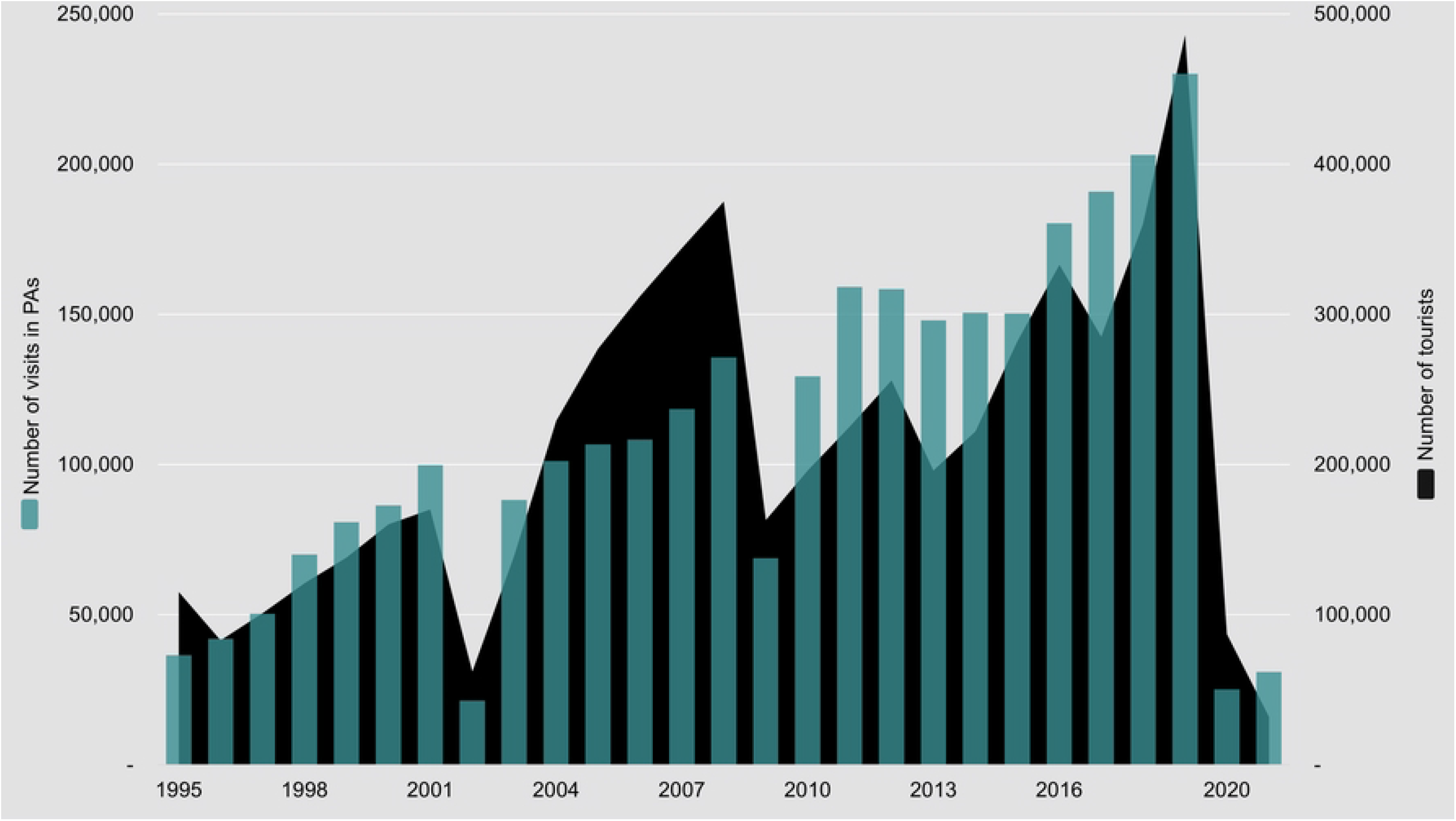
Madagascar tourists and PA entry tickets. Total number of tourists (entering Madagascar) VS. number of visits in parks and reserves (under MNP management).

The entrance fees for foreign tourists to PAs were updated and increased in 2016 and these remained unchanged until December 2021. Most PAs adopted a rate of MGA 45,000 per day for foreigners (US$ 13.38 as of 2016), with higher rates applied to the most popular terrestrial PAs and lower rates for the popular, small marine Nosy Tanihely (Fig 1, Table C in S1 File).

The closing of borders and cessation of tourism has resulted in the COVID-19 pandemic having had the most dramatic impact on numbers of entry tickets sold compared to any other crisis. The major political events in 2002 and 2009—and their aftermaths—were also cited by park managers as reasons for noteworthy decreases in ticket sales. Other significant reasons for decreases in entry ticket sales include various failures on the part of the national airline which was unable to transport tourists to remote destinations. After the 2009 coup d’état, prolonged insecurity and uncertainty led to a decrease in visitor numbers, especially to the Northern protected areas where safety and security issues also occurred. Compared with the total number of tourists in Madagascar, the total number of visits to PAs was proportionately less affected in the years following the 2002 political crisis than the years following the 2009 political crisis (Fig 4).

**Fig 4.**
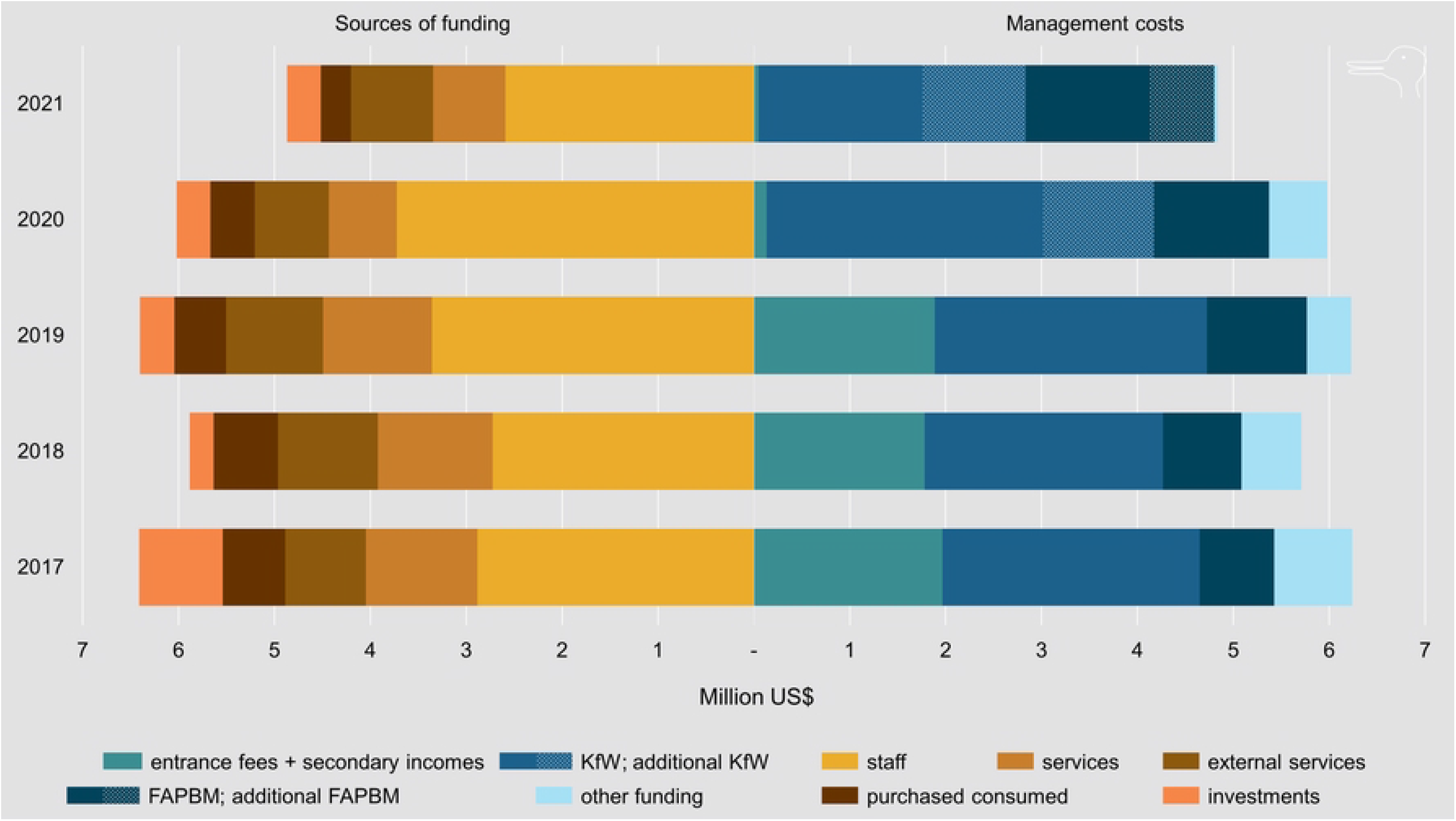
PA funding Madagascar. (MNP funding for parks and reserves in categories I, II and IV—in million US$ dollar equivalents; the losses of entrance permit ticket sales in PAs managed by MNP have been estimated to amount between US$ 3.7 and 5.9 million for 2020 and 2021—US$ 2,268,358—min = US$ 1,746,272; max = 2,855,348—in 2020 and US$ 2,415,991—min = US$ 1,878,324; max = 3,019,958—in 2021)

The FAPBM (Foundation for the Protected Areas and the Biodiversity of Madagascar) as a trust fund, is designed to provide grants to ensure the conservation of the protected areas and safeguarding of Madagascar’s biodiversity The initial capital of one million US dollars in 2005, was increased to US$ 139 million by the end of 2021. The interest generated annually is between 3 and 4 %, however, the total capital remains at a deficit when taking into account costs required to implement and maintain the high number of conservation initiatives that are currently under way in Madagascar.

Even in combination with additional income streams, the entrance fees have not generated sufficient revenue to cover the cost of the management of the network of PAs either before or after the 2016 implementation of the new, increased rates (Fig 4). MNP is predominantly supported by the German state-owned investment and development bank KfW and by FAPBM, both of which have been instrumental in assisting and dealing with the COVID-19 crisis. The emergency funds at FAPBM account for US$ 100,000 per year, but this is insufficient to address the effects of crises such as COVID-19 (Box A in S1 File). Although a new COVID-19 fund has been implemented, MNP still relies largely on finance from donors. To maintain the integrity of the PAs and support the surrounding communities, KfW granted additional funds in 2020 and 2021, as did FAPBM to compensate the communities living in the vicinity of PAs (Fig 4).

## Discussion

### The international perspective and drivers of tourism in Madagascar

Globally, the natural capital is a significant contributor to the wealth of nations [34–36]. Nature-based tourism can and does play a significant role in conservation [37,38], essentially linking intrinsic values surrounding amenity and biodiversity [39]. The pandemic has however revealed it to be a risky and a volatile sector that is vulnerable to external shocks. Financial sustainability understandably therefore, cannot be guaranteed for PAs if funding is to be solely based on revenue generated from tourist visits. As we have illustrated, PA entrance fees have never covered the cost of conservation in Madagascar despite strategic plans having been applied during the early implementation phase of the PA network in the 1990s and the National Environmental Action Plan (NEAP) during the 1990s [40].

Through implementation of the NEAP, the international donor community wanted Madagascar to create more protected areas. The government of Madagascar was willing to expand its protected area network with financial support from the international donor community. However, the promotion of PAs as the ‘goose’s golden eggs’ turned out to be an ill-founded notion. As previously outlined overall, funding remains insufficient to adequately cover the management costs of the 123 PAs. The PA network includes many extremely remote sites. Nine Strict Nature Reserve (SNR) have been reclassified as NPs, thus allowing for more development of nature-based tourism opportunities. One of those, NP Mikea has not implemented nature-based tourism following a contract with the resident populations who wish to safeguard the integrity of their park [41, but see 42]. There are other remote PAs at which nature-based tourism will not be able to be promoted for decades, because of poor infrastructure. However, such remote PAs still require funding. While entrance fees from nature-based tourism can benefit all the PAs in the network, it is clear that the financial solution lies elsewhere, even in the most successful years to date, revenue from ticket fees has not even equaled a third of the funding required to manage the network of MNP’s Pas. Additionally, during the worst year to date (2021), they dropped to a mere 1.0 % of the required revenue.

Both the protection of natural ecosystems and the benefits they provide, are optimally achieved when protected areas are supported by public policy. However, the management of protected area often suffers because of budget deficits, especially in impoverished countries where governments do not or cannot allocate resources to cover required costs [43,44]. In a recent study encapsulating almost a quarter of the world’s protected areas, more than three quarters reported a deficit of resources in staffing and budget [45]. Madagascar’s PAs were not included among the sample evaluated, but had they been, they would have increased the proportion of PAs with insufficient funding from the government.

### The local perspective on tourism in Madagascar

Madagascar’s network of PAs hosts the majority of its endemic biodiversity and intact ecosystems [28], so the provision of ecosystem services [46–48], serves to promote human well-being both actively and passively. Tourism activities conducted within and near PAs are not strictly speaking, ecosystem services, but rather, benefits delivered by nature [48].

While MNP does not manage land outside PAs, it does work closely with communities living in the external buffers of the PAs. Before 2009, MNP’s aim was to transfer half of the entrance fees generated by the parks to fund development projects benefiting the resident populations. The funds were allocated to Committees of Support to Protected Areas or COSAP, whose mission it was to identify micro-projects eligible for such funding. Villagers were grouped into associations, whose objective was to develop alternative activities for those creating pressure on the PAs. These associations would propose projects to the COSAP. Following the political crisis of 2009 and the subsequent loss of earnings from entry fees to the PAs, this redistribution was reduced, but MNP has continued to operate micro-development projects, albeit by utilising other means of funding. Additionally, MNP has systematically been advocating the creation of Local Park Committees that bring together the communities living near the PAs. MNP actively hires and trains people from these communities. It facilitates the creation of diverse occupations such as guiding for nature-based tourism, as well as the performing of miscellaneous tasks for PA management, including the organization of surveillance patrols.

It remains the intention of MNP to upgrade and maintain infrastructure in the more visited parks. Additionally, MNP has taken significant steps to improve the livelihoods of the communities residing near the parks, e.g., the building of schools and health facilities, and the developing of agricultural programmes providing viable alternatives for villagers to make their living outside of the protected areas without having to utilise resources originating from within PA boundaries. During the period of the NEAP [40], park entrance fees were equally shared with the local communities for development projects aimed at compensating people for losses of livelihood due to land reallocation for PAs—the process of which involved prohibition of access to the land and loss of the right to use the land [49].

As soon as a new PA is created, managers of the new PA are required to compensate the local communities and build capacity to enable development of the village and establish new sources of revenues. Before 2009, this was mainly achieved via the share of nature-based tourism-generated revenue, however currently, development projects are executed with special funding from FAPBM. The aim is to invest in local functions outside the PA being created to allow resident communities to obtain the products they used to harvest in the PA. Following the 2009 political crisis and various attempts to share revenues evenly with the people living adjacent to PAs, MNP reorganized its support of the local communities. Fair and equitable sharing of benefits became particularly crucial—as much as it became challenging—in a scenario where only a handful of PAs are in a position to be generating sufficient income to contribute other costs.

A significant portion of the money spent by tourists travelling to Madagascar ‘leaks’ out of the island, a common phenomenon observed in poor countries [50]. Leakage occurs when international organisations such as airlines, travel companies and hotels, pocket most of the profits. Hotels and restaurants in close proximity to the most visited PAs can earn well and do participate in the local economies, but they seldom invest in the public infrastructure required, for example that which facilitates access to the PAs. Essentially, an independent review of the entire Madagascar-related tourism value and supply chains ought to be conducted, in order to reasonably increase the benefits for the communities residing around the PAs. Included in such a review are mechanisms of sharing revenues and ways to increase local community participation [51–53].

During the period from 1995 to 2021, the entrance fees in PAs managed by MNP were increased only once at the end of 2016. With that increase of the ticket prices included, the total revenue generated from 1995 to 2021 by tourist visits to the MNP-managed PAs was less than US$ 20 million, i.e., lower than half of a percent of the revenues generated by tourism for Madagascar for these 27 years [19].

### The need for finding sustainable alternatives

Madagascar’s distinctive biodiversity is part of the global heritage. But who really pays for its conservation? [54–56]. At this point in time, when most resources are being consumed by the wealthy 10 % minority of the global population, it is patently unreasonable to expect that the often vastly poorer majority has to pay for basic ecosystem services which benefit all of humanity. As the planet’s ‘lungs’, forests are key contributors to health, stable climate, and the conservation of biodiversity [57–60]. Any economic activity should rightly therefore pay for conservation—and in so doing, for our survival within equitable structures which benefit nature. It makes sense that the international donor community covers part of the costs of the conservation of Madagascar’s biodiversity, and pledges its continued support in this regard. In order to achieve sustainability, nature-based tourism paired with grants from FABPM should cover the costs of management and of the investment in infrastructure for the entire network of PAs, as well as of a robust game plan, geared to cater for future emergency situations. Conservation governance will require alternative funding strategies if long-term financial sustainability is to be attained. With biodiversity conservation and sustainability being intertwined, we maintain that a constructive approach on the part of donors would be to consider investing funds in the PA and Biodiversity Foundation. In a world where current challenges include climate change, war, financial crises, political issues and COVID-19, the intensity of pressures on tropical forests is such that there is a real urgency for the creation and implementation of a financial support system which is an alternative to the current scenario of dependency on tourism. Only then, can harmonious balance be achieved for the sake of community well-being and biodiversity conservation.

## Acknowledgements

We would like to acknowledge the PA managers for sharing their opinion and experiences related to the fluctuation of visitors in the PAs during the sanitary crisis, especially Mrs Juliette Randriamanarivo, Director of Ambohitantely SR, Mr Mandimby Heriniaina Andriambololona, Director Ankarafantsika NP, Mr Amidou Jaovita, Director Ankarana et Analamerana SRs, Mr Mamy Tsipakay Rakotobenandrasana, Director Tsingy de Bemaraha NP, Mr Diamondra Fananako Andriambololona, Director Andranomena SR and Kirindy Mité NP, Mr Zarasolo Gérard Bakarizafy, Director Lokobe NP, Mr Solo Hervé, Head of operational section Nosy Hara and Montagne d’Ambre NPs, Mr Mora Willy Covis, Director Anjanaharibe-Sud SR and Marojejy NP, Mr Jean Fidelis Rakotomanana, Director Nosy Mangabe and Masoala NPs, Mr Clerc Tsivolany, Head of operational section Ranomafana NP, Mr Nestor Rafenonirina, Director of Sahamalaza/Îles Radama NP, Mr Justin Rakotoarimanana. Director Beza Mahafaly, SR and Tsimanampesotse and Nosy Ve Androka NPs, Mr Hery Lala Ravelomanantsoa, Director Analamazaotra and Mantadia NPs, Mrs Landisoa Randimbison, Director Nosy Tanihely NP, Mr Benoit, Director Zahamena NP, Mrs Juliette Raharivololona, Director Zombitse-Vohibasia NP, Mr Andriahery Randriamalaza, Director Mikea NP and Mr Jocelyn Bezara, Director Baie de Baly and Tsingy de Namoroka NPs. We are also grateful to colleagues at MNP, especially Mrs Haingovolatiana Raobizo Rasamoela, Ecotourism and Nature Tours Officer, Ny Saina Randrianarifetra, Legal Affairs Officer, Liliane Parany, Projects and Follow-up infractions Officer and Ghislain Rakotoniaina, Management controller, for kindly participating in the data management related to the number of visitors and financial data related of the management. We are grateful to Niamh Brannigan for inspiring early discussions. We acknowledge the comments and information provided by FAPBM, and we would also like to thank the support of the project USAID Hay Tao in Madagascar.

## Supporting Information

**S1 File. Information, data and sources considered to document Madagascar’s revenues from nature-based tourism and costs of management for conservation:**

The 43 Protected Areas considered in this study and their suitability for nature-based tourism (Table A).

Tickets sold by MNP to visit PAs (terrestrial and marine) from 2017 to 2021 (Table B).

Entrance fees rate for PAs not aligning to the rate of MGA 45, 000 starting in 2016 (Table C).

Conservation Trust Funds: The Foundation for Protected Areas and Biodiversity of Madagascar (*Fondation pour les Aires Protégées et la Biodiversité de Madagascar* FAPBM) (Box A)

